# Parallel evolution of metazoan mitochondrial proteins

**DOI:** 10.1101/047274

**Authors:** Galya V. Klink, Georgii A. Bazykin

**Affiliations:** Institute for Information Transmission Problems (Kharkevich Institute) of the Russian Academy of Sciences, Moscow 127051, Russia; Skolkovo Institute of Science and Technology, Skolkovo, 143025, Russia; Lomonosov Moscow State University, Moscow 119234, Russia

**Keywords:** fitness landscape, epistasis, parallel substitutions, heteropecilly, mitochondria

## Abstract

Amino acid propensities at amino acid sites change with time due to epistatic interactions or changing environment, affecting the probabilities of fixation of different amino acids. Such changes should lead to an increased rate of homoplasies (reversals, parallelisms, and convergences) at closely related species. Here, we reconstruct the phylogeny of twelve mitochondrial proteins from several thousand metazoan species, and measure the phylogenetic distances between branches at which either the same allele originated repeatedly due to homoplasies, or different alleles originated due to divergent substitutions. The mean phylogenetic distance between parallel substitutions is ∼20% lower than the mean phylogenetic distance between divergent substitutions, indicating that a variant fixed in a species is more likely to be deleterious in a more phylogenetically remote species, compared to a more closely related species. These findings are robust to artefacts of phylogenetic reconstruction or of pooling of sites from different conservation classes or functional groups, and imply that single-position fitness landscapes change at rates similar to rates of amino acid changes.

## Introduction

Amino acid preferences at a site, or single-position fitness landscape (SPFL, Bazykin 2015), change in the course of evolution, so that a variant conferring high fitness in one species may confer low fitness in another, either due to changes at interacting genomic sites or in the environment. These changes can be observed through phylogenetic patterns, in particular, through a non-uniform distribution of amino acid substitutions giving rise to a particular variant (homoplasies) along the phylogeny. Indeed, when a certain amino acid repeatedly arises at a particular site in a certain phylogenetic clade, but is never observed at this site in another clade, this implies that the relative fitness conferred by this variant in the former clade is higher. Different types of homoplasies – reversals, parallelisms and convergencies – have been found to be clustered on the phylogenies (Rogozin et al. 2008; Povolotskaya and Kondrashov 2010; Naumenko et al. 2012; Goldstein et al. 2015; Zou and Zhang 2015), and an attempt has been made to estimate the rate at which SPFLs change from such data (Usmanova et al. 2015).

SPFL changes in metazoan mitochondrial proteins were previously inferred from amino acid usage patterns (Breen et al. 2012), but this approach has been criticized as sensitive to the underlying assumptions regarding fitness distributions (McCandlish et al. 2013). Here, we develop an approach for the study of phylogenetic clustering of homoplasies at individual protein sites, and apply it to deep alignments of mitochondrial proteins of metazoans (Breen et al. 2012) together with their reconstructed phylogenies. Our approach compares the distributions of distances between parallel and divergent substitutions to infer robustly changes in relative fitness of different variants at a site between branches of the phylogenetic tree.

## Materials and Methods

### Alignment and phylogeny

We obtained multiple-species alignments of 12 mitochondrial proteins of metazoans from (Breen et al. 2012), and analyzed alignment columns with fewer than 1% gaps (which comprised 77% of all sites). As there is no accepted phylogenetic tree for this large and diverse set of species spanning a wide range of phylogenetic distances, we took a hybrid approach to reconstructing their phylogeny. First, we constrained the tree topology using the curated taxonomy-based phylogeny of the ITOL (Interactive tree of life) project (Letunic and Bork 2007). By requiring the presence of each species in the ITOL database, we were left with >900 metazoan species for each protein (Table 1). The resulting topology was not fully resolved, and contained multifurcations. We then used RAxML 8.0.0 (Stamatakis 2014) under the GTR-Γ model to resolve multifurcations and to estimate the branch lengths. Finally, ancestral states were reconstructed using codeml program of the PAML package (Yang 1997) under the substitution matrix and the value of the parameter alpha of the gamma distribution inferred by RAxML.

**Table 1.**
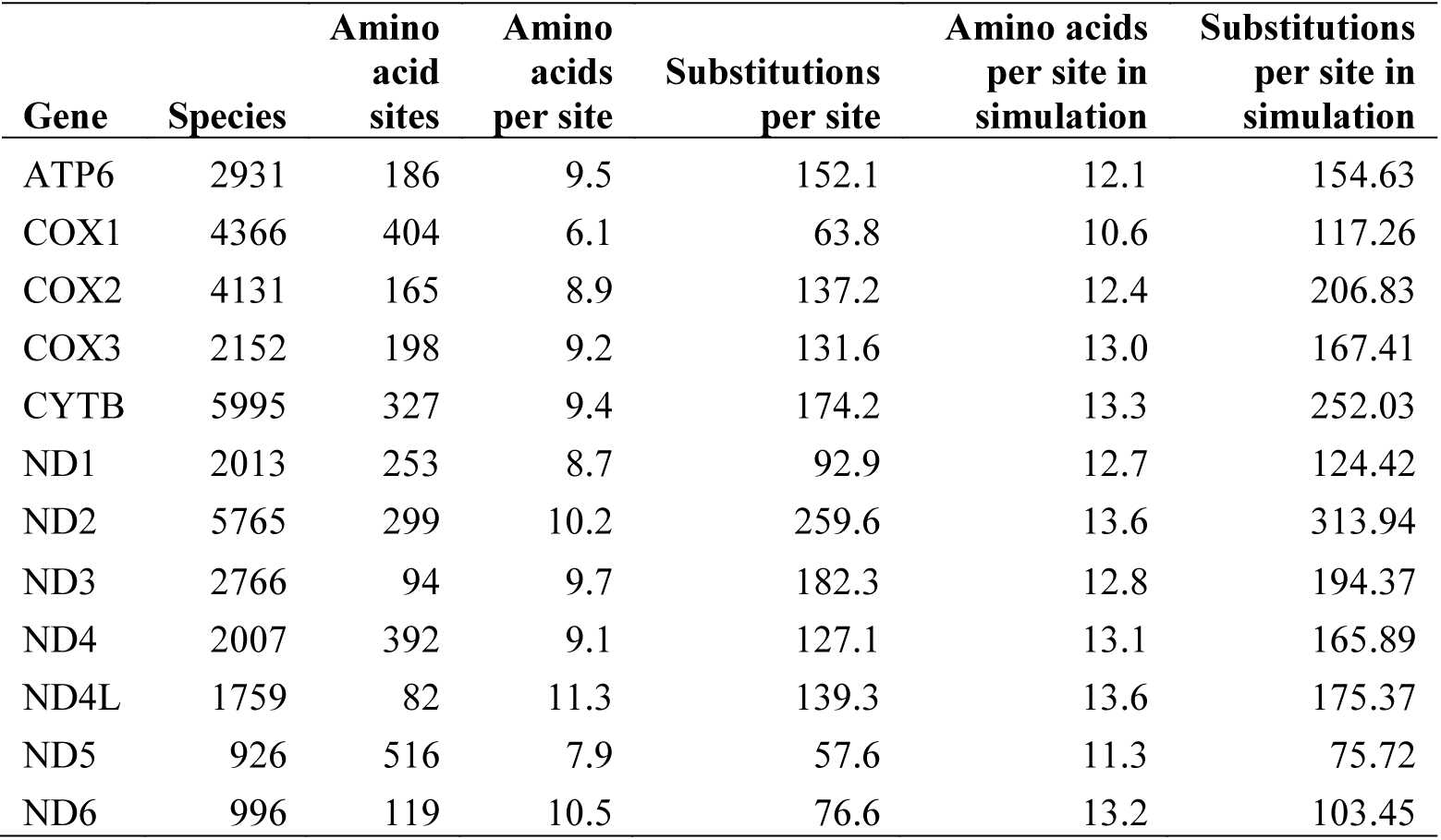
Amino acid substitutions in mitochondrial genes of metazoans.

Independently, using the same methods and parameters, we reconstructed a joint phylogeny of 3586 chordate, 586 non-chordate and 178 fungal species (for a total of 4350 species) based on amino acid sequences of five concatenated mitochondrial genes, covering a total of 1524 amino acid positions.

Transmembrane and non-membrane sites were obtained from UniProt database (http://www.uniprot.org/).

### Clustering of substitutions on a phylogeny

Using the inferred states of amino acid sites at each node, we inferred the positions of all substitutions at all protein sites on the phylogenies of the corresponding proteins. For each ancestral amino acid at a site, we defined parallel substitutions as those giving rise to the same derived amino acid, and divergent substitutions, as those giving rise to different derived amino acids (fig. 1A). We considered only those pairs of substitutions that happened in phylogenetically independent branches, i.e., such that one was not ancestral to the other. For subsequent analyses, we used only homoplasy-informative sites, i.e., sites that have at least one pair of parallel substitutions and one pair of divergent substitutions from the same ancestral amino acid. The phylogenetic distance between a pair of substitutions was defined as the distance (measured in the number of amino acid substitutions per amino acid site inferred by RAxML) between the centers of the edges where those substitutions have occurred, i.e., the sum of the distances from the centers of these edges to the last common ancestor of the two substitutions (fig. 1A). Alternatively, as a proxy for the site-specific evolutionary distance, we multiplied the phylogenetic distance by the codeml estimate of the site-specific substitution rate.

**Figure 1.**
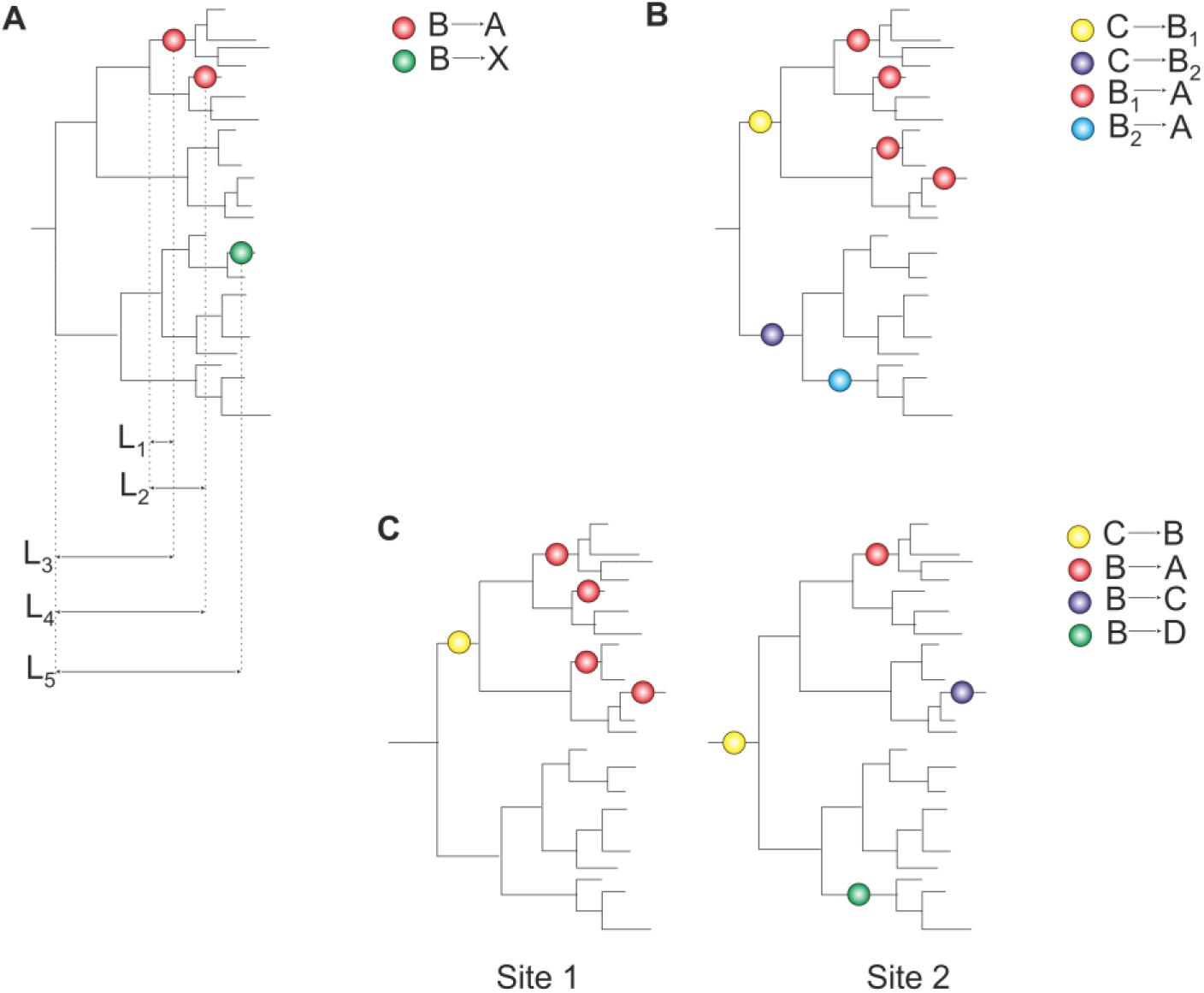
Inference of phylogenetic distances between parallel and divergent substitutions. Dots represent substitutions mapped to nodes of a phylogenetic tree. (A) For each pair of amino acids (B, A) at a particular amino acid site, we consider the distances between all parallel B→A substitutions (L_1_+L_2_), and distances between all divergent substitutions B→A and B→X (L_3_+L_5_, L_4_+L_5_), where X is any amino acid other than A and B. (B) The B_1_→A substitution is more frequent than the B_2_→A substitution, leading to an excess of homoplasies at small phylogenetic distances when parallel and convergent substitutions are pooled together. (C) Pooling of sites with different properties may also lead to an excess of homoplasies at small phylogenetic distances (see text).

To compare the distances between parallel and divergent substitutions while circumventing the potential biases associated with pooling sites and amino acids with different properties (see below), we subsampled the pairs of parallel and divergent substitutions, and analyzed the distances in this subset. For this, for each ancestral amino acid at homoplasy-informative sites, we picked randomly min(*N*_paral_, *N*_diverg_) pairs of parallel substitutions and the same number of pairs of divergent substitutions, where *N*_paral_ and *N*_diverg_ are the numbers of parallel and divergent substitutions originating from this amino acid. We repeated this procedure for all ancestral amino acids at all sites, thus obtaining two equal-sized subsamples of parallel and divergent substitutions, and measured all distances in these resulting subsamples. The parallel to divergent ratio (P/D) for each distance bin was calculated by dividing the number of parallel pairs of substitutions by the number of divergent pairs of substitutions that had occurred at a distance from each other falling into this bin. This statistic is closely related to the O-ring statistic widely used in spatial ecology to measure aggregation in communities (Wiegand and A. Moloney 2004).

To obtain mean values and confidence intervals of each statistic, we bootstrapped sites in 1000 replicates, each time repeating the entire resampling procedure.

### Robustness of tree shape and ancestral states reconstruction

We identified a set of high-confidence pairs of substitutions, defined as follows. For each branch of the phylogenetic tree, we obtained the bootstrap support value in 100 bootstrap replicates using RAxML. A pair of substitutions was considered high-confidence when (i) at least one node between substitutions had 100% bootstrap support, ensuring the robustness of these nodes; and (ii) for each substitution from the pair, the maximum likelihood estimate for amino acids in ancestral and derived nodes was equal to 1, ensuring the robustness of ancestral state reconstruction.

### Simulated evolution

For each gene, we simulated amino acid evolution using the evolver program of the PAML package (Yang 1997), under the empirical_F model and discrete-gamma distributed rates between sites. The phylogenetic tree, substitution matrix, alpha parameter and number of categories for discrete gamma of the gamma-distribution were obtained from the output of RAxML for the corresponding gene. From the amino acids thus simulated for the leaves of the tree (i.e., extant species), we then reconstructed the ancestral states using codeml under the same parameters.

### Simulated evolution under different substitution matrices

To test the effect of differences in substitution matrices between clades due to clade-specific biases, we obtained individual RAxML-generated matrices for each of the three major groups of species in the joint 5-gene phylogeny: chordates, non-chordates and fungi, using the same methods as above. We then used these matrices to simulate evolution of the corresponding groups of species of the joint 4350-species tree, and used generated sequences for the analysis.

## Results

### Phylogenetic clustering as evidence for SPFL changes

We devised an approach for analysis of the clustering of parallel substitutions at a site. While conceptually related to the previous methods (Povolotskaya and Kondrashov 2010; Goldstein et al. 2015; Zou and Zhang 2015), it is designed to be robust to other specifics of the phylogenetic distribution of substitutions, and to control for any potential biases that can arise from pooling sites with different properties. Since it is difficult to obtain robust evidence for SPFL changes for an individual amino acid site even using large numbers of species, getting a significant signal of SPFL changes requires pooling different amino acid sites. The problem is that these sites may differ in their properties, and such differences may lead to artefactual evidence for SPFL changes, for the following reasons.

First, pooling of parallel and convergent substitutions giving rise to the same descendant variant, i.e., substitutions with the same and different ancestral variants, may provide artefactual evidence for SPFL changes due to reasons such as the structure of the genetic code. For example (fig. 1B), an amino acid A within a clade may arise repeatedly from the ancestral amino acid B_1_ at a particular clade simply because the B_1_→A mutation is frequent. If another amino acid B_2_ is more prevalent than B_1_ at a different clade, and the B_2_→A mutation is less frequent, this will lead to an excess of substitutions giving rise to A in the former clade; this excess, however, is not an evidence for SPFL changes, but instead occurs for non-selective reasons. To control for this, we do not consider convergent substitutions, and separately consider the distributions of parallel and divergent substitutions from each ancestral variant B.

Second, even independent consideration of different ancestral variants still permits clustering of homoplasies without SPFL changes when sites, and amino acids within sites, with diverse properties are pooled together. To illustrate this, assume that we analyze phylogenetic distances between parallel and divergent substitutions in a pooled sample of sites. Consider the hypothetical scenario in Figure 1C. At site 1, the amino acid B only resides within a relatively small clade. Therefore, both B→A and B→X substitutions are, by necessity, phylogenetically close to each other. By contrast, at site 2, the amino acid B is long living, and the distances between substitutions from it may be larger. If such sites also differ systematically in their amino acid propensities, this might lead to artefactual evidence for SPFL changes. For example, if sites where B is short-living (like site 1) also tend to be those where few amino acids confer high fitness (so that B→A substitutions are more frequent), while sites where B spans a large clade tend to be promiscuous with respect to the occupied amino acid (so that B→X substitutions are more frequent), pooling such sites may result in an excess of homoplasies within short phylogenetic distances.

We circumvent this problem by resampling matched sets of parallel and divergent substitutions. Specifically, at each homoplasy-informative site (see Methods), we consider different ancestral amino acids separately. For each ancestral amino acid B, we subsample our sets of pairs of substitutions: for each pair of parallel (B→A, B→A) substitutions, we randomly pick exactly one pair of divergent (B→A, B→X) substitutions from the same site. Finally, we pool together these subsamples from different sites, and analyze distances between parallel and divergent substitutions in this pooled set. This approach controls for any possible biases associated with differences in phylogenetic distributions of different ancestral amino acids. From the resulting subsets of distances, we calculate the ratio of the numbers of parallel to divergent substitutions (P/D) for each distance bin (see Methods).

### Parallel substitutions in mitochondrial proteins are phylogenetically clustered

We applied this approach to the phylogenetic trees of 12 orthologous mitochondrial proteins of metazoans, each including >900 species. At the vast majority of sites, we observe many amino acid variants, in line with (Breen et al. 2012). By reconstructing the ancestral states and substitutions at each site, we observe that most of these variants have originated more than once, allowing us to study the phylogenetic distribution of homoplasies in detail (Table 1 and supplementary table 1, Supplementary Material online).

We observe an excess of parallel substitutions for species at small phylogenetic distances from each other (fig. 2-3), in line with the previous findings in vertebrates that used a smaller dataset (Goldstein et al. 2015). The P/D ratio is ∼1.7 to 2.5 at phylogenetic distances less than 0.1, but drops to ∼1 rapidly for larger distances (fig. 4 and supplementary fig. 1, Supplementary Material online). In simulated data, only a very weak decrease in the P/D ratio was observed, which is possibly attributable to minor biases in phylogenetic reconstruction.

**Figure 2.**
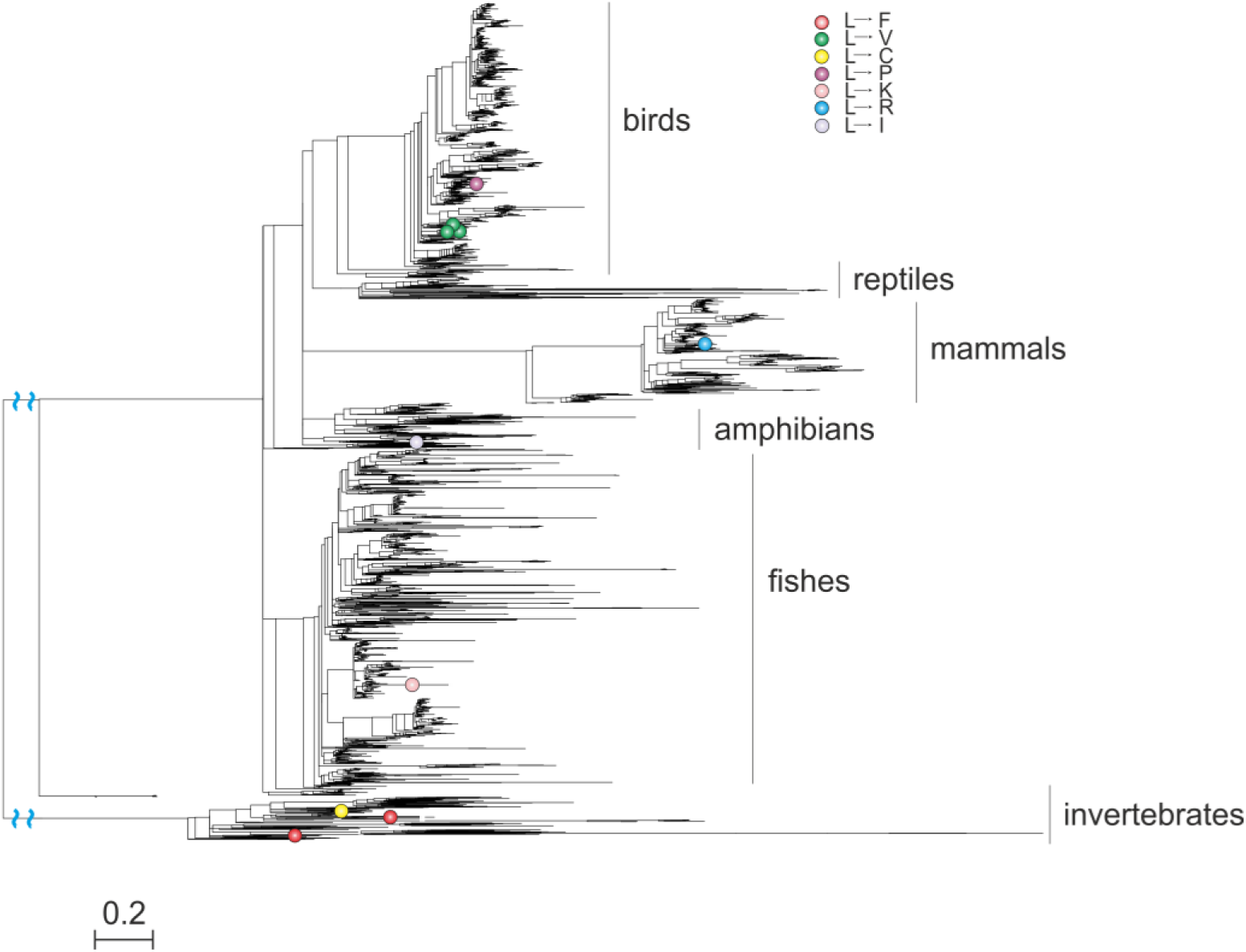
Parallel and divergent substitutions at site 202 of ATP6 (NCBI reference sequence numbering for the human sequence). The ancestral variant (L) has experienced multiple substitutions, which are scattered throughout the phylogeny. However, the two parallel L→F substitutions occur in closely related species; the same is true for the three parallel L→V substitutions. Phylogenetic distances are in numbers of amino acid substitutions per site. The branches indicated with the blue waves are shortened by 1.2 distance units.

**Figure 3.**
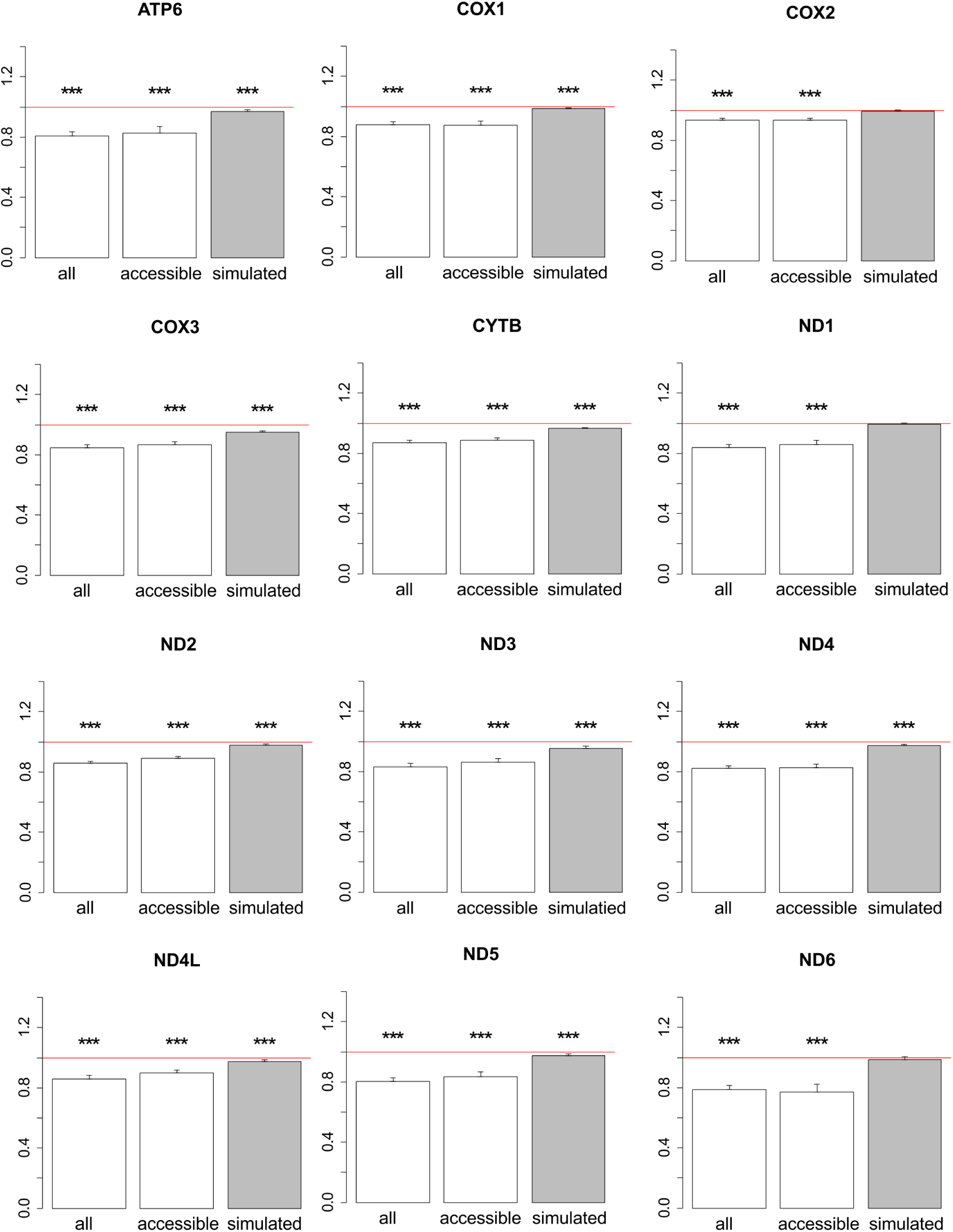
Ratios of phylogenetic distances between parallel and divergent substitutions in metazoan phylogenies. Values below 1 imply that the parallel substitutions are closer at the phylogeny to each other, compared to divergent substitutions. The bar height and the error bars represent respectively the median and the 95% confidence intervals obtained from 1,000 bootstrap replicates, and asterisks show the significance of difference from the one-to-one ratio (red line; ***, P<0.001; no asterisk, p>0.05). all, real data; accessible, real data only for substitutions from accessible amino acid pairs (see text); simulated, simulated data.

**Figure 4.**
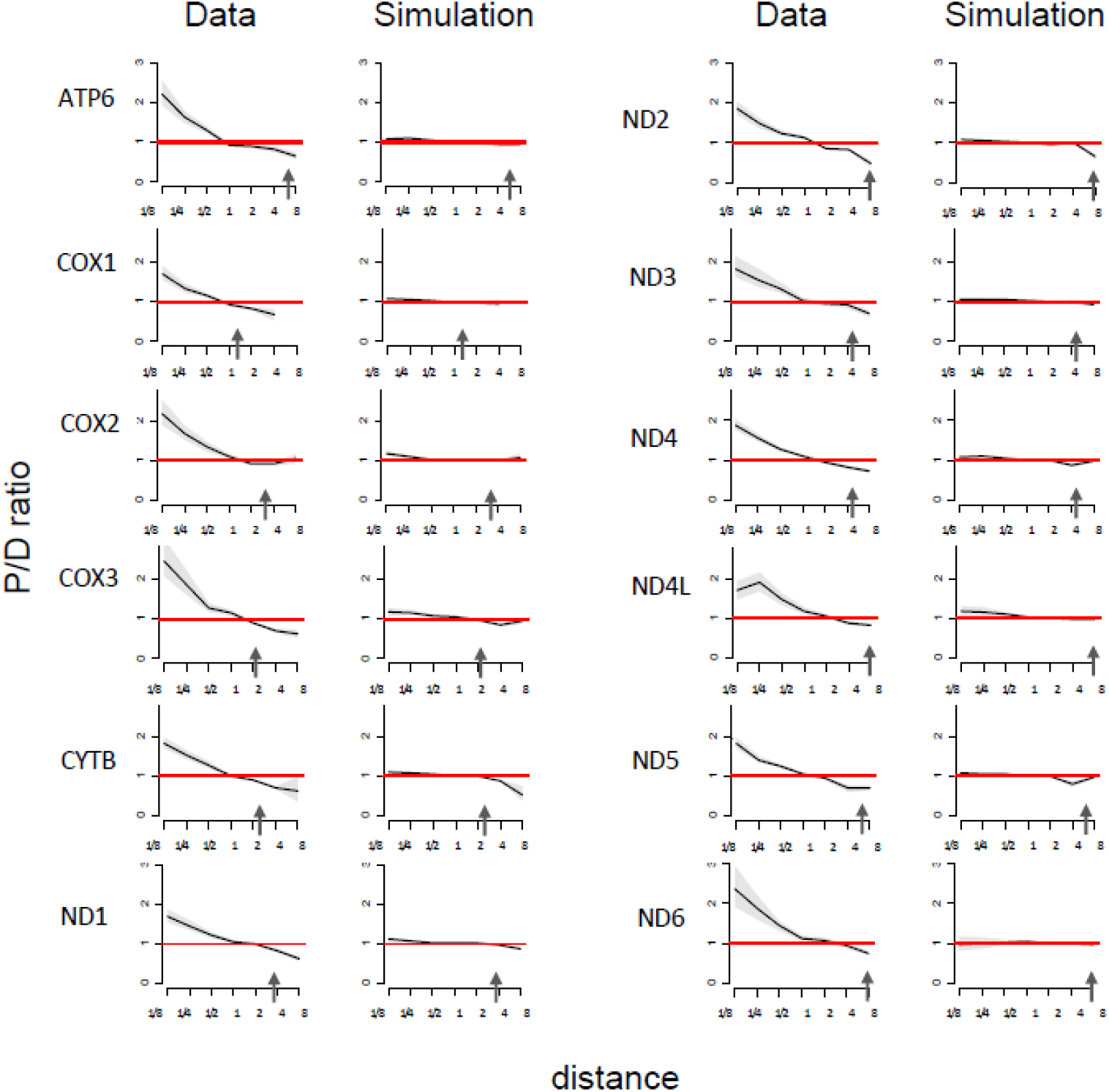
Higher fraction of parallel substitutions between closely related species. Horizontal axis, distance between branches carrying the substitutions, measured in numbers of amino acid substitutions per site (split into bins by log_2_(distance)). Vertical axis, P/D ratios for substitutions at this distance. Black line, mean; grey confidence band, 95% confidence interval obtained from 1000 bootstrapping replicates. The red line shows the expected P/D ratio of 1. Arrows represent the distance between human and *Drosophila*.

**Figure 5.**
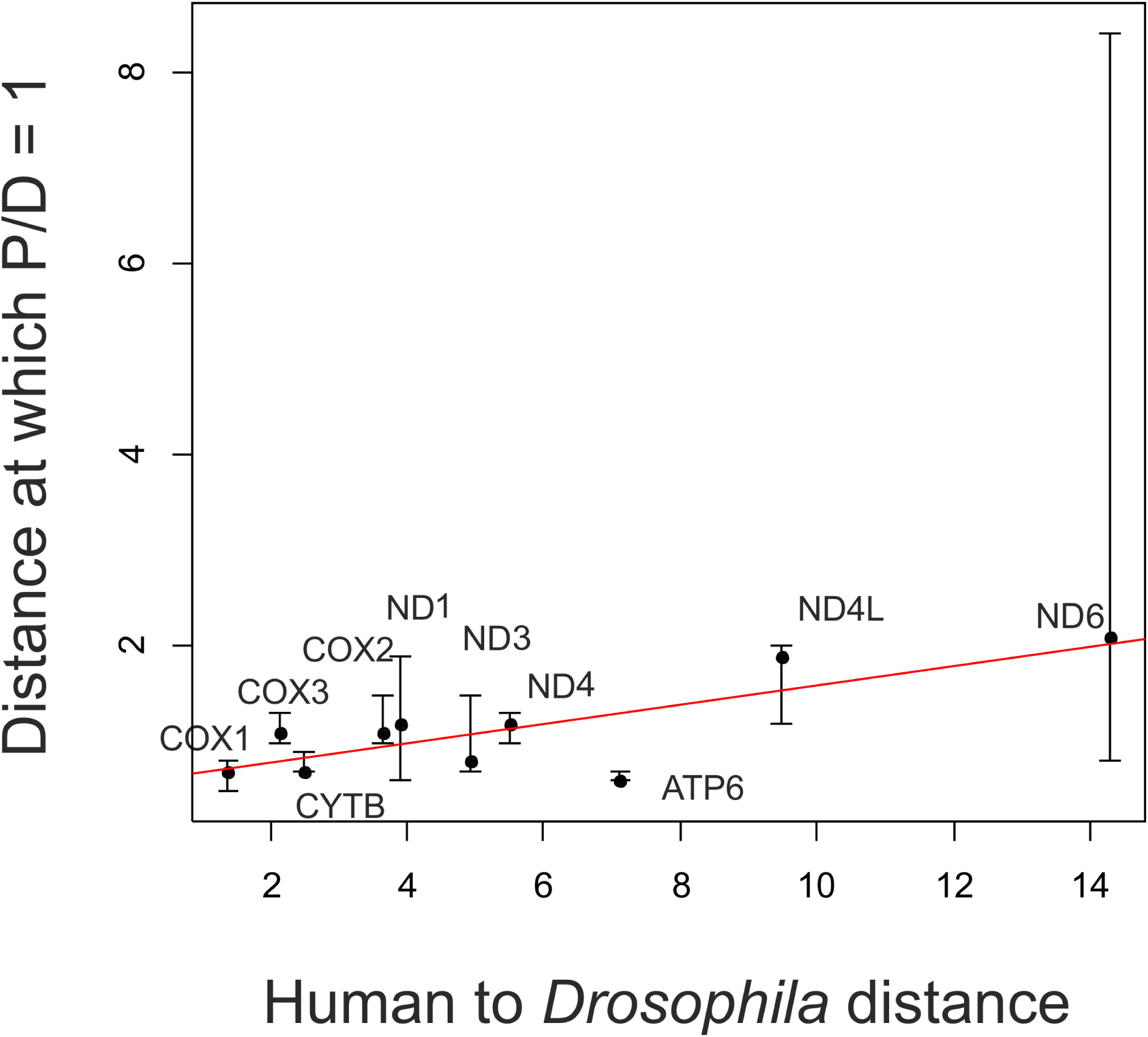
Faster decline of the P/D ratio for rapidly evolving genes. Horizontal axis, gene-specific phylogenetic distance between *Homo sapiens* and *Drosophila simulans*. Vertical axis, phylogenetic distance at which the P/D ratio reaches 1. Each dot represents one gene, and the line represents the linear trend. Only the ten genes with available *Drosophila simulans* sequences were used. Error bars correspond to the 95% confidence interval for the distance at which the P/D ratio reaches 1, obtained by bootstrapping sites 1000 times.

We also measured P/D ratios for phylogenetic distances normalized by site-specific evolutionary rates (see Methods) and obtained similar plots (Supplementary fig. 2, Supplementary Material online). We also asked whether the mean P/D distance is different between sites with different evolutionary rates, but saw no systematic differences (Supplementary fig. 3, Supplementary Material online). These findings suggest that the P/D ratios are more sensitive to the evolutionary distance spanned by the species rather than by the individual site. Finally, we observed similar effect in trans-membrane and in nonmembrane residues. In most proteins, the effect is slightly weaker in transmembrane residues, but this difference is extremely weak (Supplementary fig. 4, Supplementary Material online).

The rate at which the P/D ratio declines with phylogenetic distance varies a lot between genes. We asked whether this difference has to do with the intrinsic rate of protein evolution, which also varied strongly between genes. We used the number of substitutions between human and *Drosophila* as the proxy for the evolutionary rate of the protein, with higher values corresponding to rapidly evolving genes; and the phylogenetic distance at which the P/D ratio (which is initially always larger than one) reaches one, as the proxy for the rate of the decline of the P/D ratio, with higher values corresponding to a slower decline. Among the 10 analyzed genes for which sequences both for human and *Drosophila* were available, the P/D decline appeared to be somewhat faster in fast-evolving genes, although this trend was not significant (Spearman’s test: R=0.53, p=0.11).

### Excess of parallel substitutions at small phylogenetic distances is not an artefact

Conceivably, the decline of the P/D ratio could be an artefact of erroneous phylogenetic reconstruction. Indeed, if a clade is erroneously split on a phylogeny, synapomorphies (shared derived character states) may be mistaken for parallel substitutions, and this is more likely for closely related species (Mendes et al. 2016). We tested the contribution of such artefacts by performing our analyses only on high-confidence pairs of substitutions (see Methods). Our definition of this set was conservative, because branches with parallel substitutions are expected to have a reduced bootstrap support, as such substitutions cause attraction of the branches where they occur in phylogenetic reconstruction. Indeed, in the Breen et al. dataset, this procedure discarded the vast majority of parallel substitutions at very small phylogenetic distances, leaving us with too little data. To circumvent this, we reconstructed a 5-gene, 4350-species phylogeny (see Materials and Methods for details). Similarly to the main analysis, the P/D ratio declined monotonically with distance between substitutions in this dataset (Figure 6), implying that the excess of homoplasies at short phylogenetic distances is unlikely to be an artefact of phylogenetic reconstruction.

**Figure 6.**
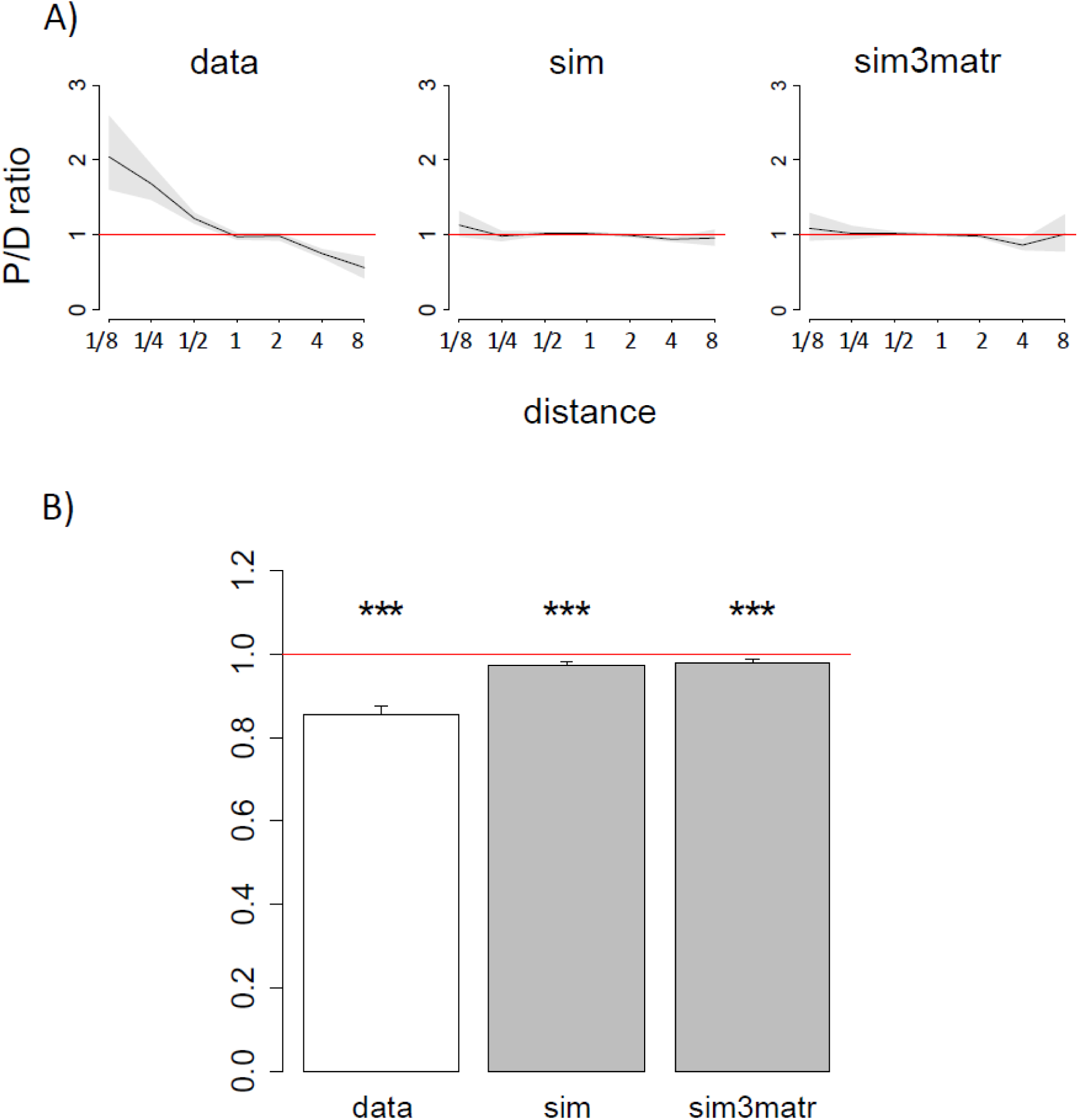
Higher fraction of parallel substitutions between closely related species (A) and ratios of phylogenetic distances between parallel and divergent substitutions (B) in the 4350-species phylogeny, for high-confidence pairs of substitutions. sim3matr, simulation with independent substitution matrices for each clade (see text).

Changes in the P/D ratio with increasing phylogenetic distance imply changes in the rate of the B→A substitution relative to other substitutions. The rate of a substitution is the product of the mutation and fixation probabilities, and changes in the substitution rate may arise from differences in either of these processes between clades.

Can the changes in substitution rates with phylogenetic distance be explained by changes in mutation rates? There can be two scenarios for this. First, the mitochondrial mutational spectra could differ between clades, potentially leading to differences in the rate at which a particular mutation occurs. If the mutation rate corresponding to a particular substitution is much higher in a particular clade, compared to the rest of the phylogeny, this may lead to an excess of homoplasies falling into this clade. To ask whether this mechanism contributes, we simulated evolution of the three major clades of the 4350-species phylogeny using an independent substitution matrix for each clade (see Materials and Methods for details) and performed our analysis for the whole phylogeny using simulated sequences. Results for this simulation are indistinguishable from the simulation constructed with one matrix for all clades (Fig. 6), implying that this mechanism is unlikely to cause the observed pattern. Moreover, most of the change in the P/D ratio occurs at very small phylogenetic distances (fig. 4), where the mutation matrices are very similar, and unlikely to contribute to our effect.

Second, even if the changes in the P/D ratio are not due to changes in the overall mutation matrix, they may still arise from differences in codon usage. This may be observed if amino acid B tends to be encoded by different codons in the two clades, and the rate of the parallel B→A substitution is higher in the clade where B is encoded by a codon that predisposes to this mutation. To test this, we defined “accessible” amino acid pairs as those (B, A) pairs where A can be reached through a single nucleotide substitution from any B codon, and considered such accessible pairs independently. In this subset, the excess of parallel changes at small phylogenetic distances was also observed (fig. 3), which means that it is not caused by the structure of the genetic code.

Finally, we analyzed the distribution of the pairs across the phylogenetic tree. Both in data and in simulations, substitutions with high (or low) P/D are clustered on the tree. The distribution of parallel (as well as divergent) substitutions over the tree is non-uniform mainly because of differences in branch lengths: longer branches carry more substitutions of all kinds. To test if there is an excess of branches with more than expected parallel substitutions, we plotted the distribution of pairs of branches by the number of parallel substitutions that had occurred in them (Supplementary figure 5). We see no systematic differences between the distributions obtained from the data and from simulations, implying that our results are not driven by clustering of substitutions in some of the branch pairs. Moreover, in branch pairs that carried many substitution pairs in them, both branches tended to be long, leading to large distances between the two substitutions in a pair; while most of the effect is observed at small distances (Figure 4).

For three of the genes, we also randomly picked and examined visually 20 pairs of parallel and 20 pairs of divergent substitutions among those with distances <0.1 between them. Parallel and divergent pairs in data and in simulations were clustered in the same parts of the tree, demonstrating that this effect is likely to be determined by the shape of the tree. Specifically, they were clustered in the regions of the tree with many short branches, which are the only ones contributing to the phylogenetically close pairs of substitutions in Figure 4.

## Discussion

The rate at which a specific substitution occurs is a monotonic function of fitness differences between the descendant and the ancestral variants, and changes in the substitution rates in the course of evolution indicate that these fitness differences, and thus the SPFL, change. Obtaining the entire substitution matrix for individual amino acid sites is problematic even given hundreds of species (Rodrigue 2013); still, the changes in the relative substitution rates can be inferred using some summary statistics. The change in the extent of parallelism in the course of evolution is one such convenient statistic: as a substitution becomes more deleterious, its rate decreases, and it becomes more sparsely distributed on the phylogeny.

We observe that parallel substitutions in the evolution of metazoan mitochondrial proteins are phylogenetically clustered; i.e., that such substitutions are more likely to occur in the phylogenetic vicinity of each other, compared with divergent substitutions. As a result, the distance between two parallel substitutions on the phylogenetic tree is, on average, ∼20% lower than the distance between divergent substitutions, or than expected if their rate was constant across the tree (fig. 3). We show that these results are unlikely to be artefacts of phylogenetic reconstruction or of pooling together sites and amino acids with different properties. Our results cannot be explained by a simple covarion model, in which a site alternates between neutral and constrained (Fitch and Markowitz 1970; Fitch 1971), as the changes we observe are not associated with changes in the overall substitution rates. For the same reason, they also cannot be explained by a broader class of heterotachy models in which the overall rates of evolution of a site vary with time (Lopez et al. 2002; Yang and Nielsen 2002; Murrell et al. 2012), but require heteropecilly (Tamuri et al. 2009; Roure and Philippe 2011), i.e., variation with time of rates of individual substitutions. We show that these differences are unlikely to be caused by systematic gene-or genome-wide differences in substitution matrices between clades, which may result from differences in mutation patterns, selection for nucleotide or amino acid usage, or gene conversion.

Instead, they most likely reflect changes in single-position fitness landscapes (Mustonen and Lässig 2007; Bazykin 2015) that accumulate in the course of evolution. Indeed, site-specific differences in the rate of a substitution leading to a particular amino acid imply that the relative fitness of this amino acid relative to other amino acids at this site changes with time. Decrease in this frequency with phylogenetic distance may be caused by a decline in the fitness of this allele, and/or by an increase in the fitness of other alleles; it is hard to distinguish between these possibilities with the available data, although both factors likely play a role (Naumenko et al. 2012). Our single-site approach also prevents us from distinguishing between the possible causes of the SPFL changes: evolution of other sites (of the same or other proteins) involved in epistatic interactions with the focal site, environmental changes, or perhaps a combination of both.

The numbers of substitutions to the same or to another amino acid, i.e. convergent and divergent substitutions at different phylogenetic distances, have been used previously to characterize evolution. In ancient proteins, the rate of convergence monotonically decreases with phylogenetic distance, and half of the reversals were estimated to become forbidden after 10% protein divergence (Povolotskaya and Kondrashov 2010). The ratio of the rates of convergent and divergent substitutions drops by more than twofold with phylogenetic distance within vertebrates (Goldstein et al. 2015). Similarly, the rate of convergence decreases with phylogenetic distance in mammals and fruit flies (Zou and Zhang 2015). Our analysis spans larger phylogenetic distances than that of Goldstein et al. (2015); still, most of the observed effect is local (fig. 4). On the basis of the data from different sources, and assuming a two-state fitness space such that each amino acid variant at a particular amino acid site can be either “prohibited” or “permitted”, the rate at which a particular variant switches between these two states has been estimated as ˜5 such switches per unit time required for a single amino acid substitution to occur at this site (Usmanova et al. 2015). In our data, the rate of SPFL change appears to vary widely between proteins, as the time necessary for the P/D ratio to reach 1 varies between 0.6 for ATP6 and 2.1 for ND6. It also is strongly dependent on the size and the shape of the phylogeny. Still, in our data, the rate of SPFL changes has roughly the same scale as the rate of amino acid evolution (fig. 4), which is consistent with the results of (Mustonen and Lässig 2007) who have shown that fluctuations in SPFLs of *Drosophila* proteins occur with rates comparable with neutral mutation rates.

In summary, our results allow to suggest that the fitness landscapes of amino acid sites of mitochondrial proteins change with time, supporting previous conjectures that such landscapes are dynamic in this dataset (Breen et al. 2012; Breen et al. 2013). Whether these changes are driven by changes in the intra-protein or inter-protein genomic context between species, or by environmental changes, remains a subject for future research.

## Acknowledgements

This work was supported by the Russian Science Foundation grant no.14-50-00150.

We thank Shamil Sunyaev, Alexey Kondrashov, Alexander Favorov, Andrey Mironov, Dmitry Pervouchine, Vladimir Seplyarskiy, Sergey Naumenko, Elena Nabieva, Ivan Cytovich, David McCandlish, Joshua Plotkin, Richard Goldstein and Dinara Usmanova for valuable comments.

---

**Supplementary figure 1.** Numbers of pairs of parallel (P) and divergent (D) substitutions. For each distance window of size 0.1, the two ends of the bar correspond to the average (across 1000 bootstrap replicates) numbers of P and D, so that bar length is equal to the absolute value of their difference. Red bars correspond to P>D (so that the top of the bar corresponds to P, and the bottom of the bar, to D), and blue bars correspond to P<D (so that the top of the bar corresponds to D, and the bottom of the bar, to P).

**Supplementary figure 2.** Higher fraction of parallel substitutions between closely related species in metazoan phylogenies. Horizontal axis, distance between branches carrying the substitutions, measured in numbers of amino acid substitutions per site (log_2_distance bins). Vertical axis, P/D ratios for substitutions at distances falling into this distance bin. Black line, mean; grey confidence band, 95% confidence interval obtained from 1000 bootstrapping trials. The red line shows the expected P/D ratio of 1. The figure is similar to Figure 4, but phylogenetic distances between substitutions were measured using branch lengths normalized by site-specific evolutionary rates obtained by codeml.

**Supplementary figure 3.** Ratios of phylogenetic distances between parallel and divergent substitutions in metazoan phylogenies, for sites from different rate categories (1: slowest-evolving sites; 4: fastest-evolving sites). Notations are the same as in Figure 3.

**Supplementary figure 4.** Ratios of phylogenetic distances between parallel and divergent substitutions in metazoan phylogenies, for transmembrane (mem) and non-membrane (non-mem) sites. Red asterisks represent p-values of the 2-sided Wilcoxon rank sum test with the null-hypothesis of no difference between two types of sites (***, P<0.001), with signs > or < indicating the direction of the difference. Other notations are as in Figure 3.

**Supplementary figure 5.** The distribution of pairs of branches by the fraction of all parallel substitutions that had occurred on them. Red, data; blue, simulation.

